# RNAi-mediated HBV antigen shutdown enhances the antiviral immune effects of PEGIFNα via altering T and B cell crosstalk

**DOI:** 10.1101/2024.12.03.626539

**Authors:** Wenjing Zai, Kongying Hu, Mengying He, Ziyang Song, Chen Luo, Minxiang Xie, Asha Ashuo, Jieliang Chen, Zhenghong Yuan

## Abstract

PEGylated interferon-α (PEGIFNα) demonstrates promising therapeutic outcomes against chronic hepatitis B (CHB), whereas patient response to PEGIFNα therapy remains unsatisfied. Shutdown of hepatitis B virus (HBV) antigens by RNA interference (RNAi) could enhance PEGIFNα efficacy in CHB patients, whereas the underlying immunological mechanisms remain obscure. We performed studies by utilizing our newly established extracellular humanized IFNAR (IFNAR-hEC) mice. An in-house constructed small interfering RNAs (GalNac-siHBV) was administrated to mice either alone or in combination with PEGIFNα. The phenotypic and functional characteristics of peripheral and organ-specific immune cells were assessed by flow cytometry, ELISpot, RNA sequencing (RNA-seq), and single-cell RNA-seq (scRNA-seq) analysis. Our results demonstrated that combined treatment with PEGIFNα and RNAi exerted a synergistic and prolonged inhibition of HBsAg (∼4log_10_ IU/mL, vs PBS) and induced a higher incidence of HBsAg seroconversion (∼30%), comparing with either monotreatment. Mechanistically, combined therapy improved the functionality of global T and B cells, triggered increased anti-HBs producing B cells, and enhanced IFNγ-producing T cells. scRNA-seq analysis revealed that the combined therapy reduced inhibitory B cell-B cell interaction, enhanced MHC-I signaling mediated T cell-T cell communication, and improved T cell-B cell crosstalk, thus improving the functionality of T and B cells. Enhanced MHC-II signaling networks across B cells and hepatocytes/Cd8^+^ T cells further promoted HBsAg seroconversion in the combined treatment groups. These results together provided scientific rationale and lessons for the combination of the two towards better therapeutic efficacy.

**Graphical Abstract:** 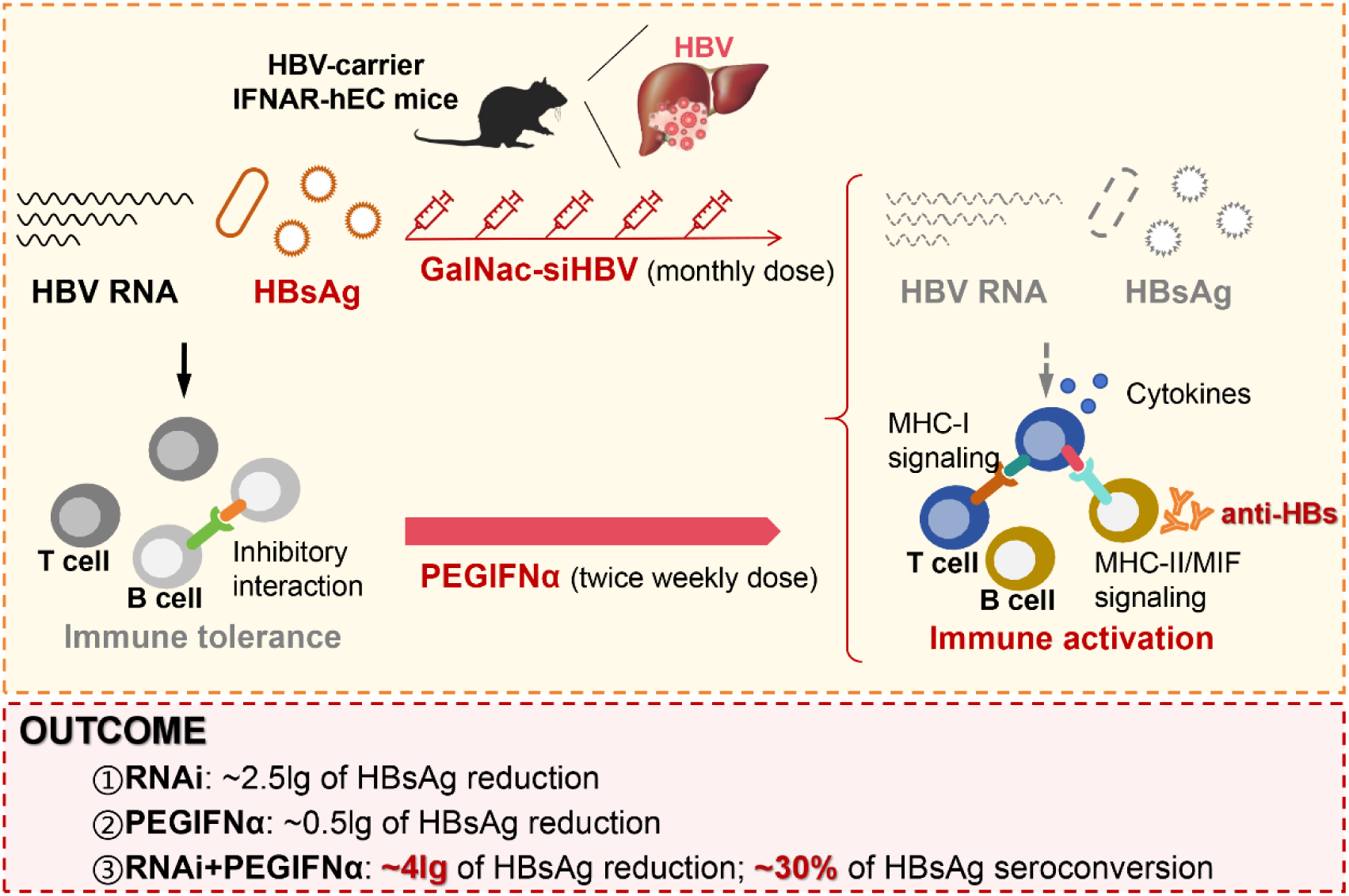

**Highlights:**

1. Shutting down HBsAg through RNA interference augmented the antiviral immune effects of PEGIFNα in chronic HBV-carrier IFNAR-hEC mice.
2. Combined RNAi plus PEGIFNα augmented the functionality of T cells, promoted B cell activation and class switch, but also exerted some suppressive effects on B cells.
3. Reduced inhibitory B cell-B cell interaction, enhanced MHC-I signaling between T cells and T cells, and improved T cell-B cell crosstalk, improved the functionality of T cells and B cells.
4. Enhanced MHC-II signaling networks across B cells and hepatocytes/ Cd8^+^ T cells further promoted HBsAg seroconversion in RNAi plus PEGIFNα combined treatment groups.

## INTRODUCTION

Over 292 million people worldwide are suffering chronic hepatitis B (CHB),^1^ which is a major risk that leads to liver fibrosis, cirrhosis and hepatocellular carcinoma, and current therapeutic options are limited.^2^ The persistence of covalently closed circular DNA (cccDNA) and integration of HBV DNA into the host genome together makes the goal of “complete cure” unachievable.^3^ The “functional cure” of CHB, which is characterized by the clearance of seral HBsAg and HBV DNA, is still a challenging endpoint.^4^ Nucleos(t)ide analogues (NUCs) are the most widely used clinical therapies that reduce HBV replication, but HBsAg loss is rarely observed even after decades of therapy.^5^ PEGylated type I interferons (PEGIFNα) possess both antiviral and immunomodulatory effects, whereas the efficacy is moderate, and sustained control of the virus is rare to meet.^6,7^ Besides, repeated PEGIFNα treatment can cause systemic toxicity and immunosuppressive effects.^8^ Recent studies showed that PEGIFNα alone does not improve peripheral HBV-specific CD8^+^ T cell responses even upon successful treatment.^9^ No significant improvement of T cell proliferation, cytokine production or cytotoxicity was observed during PEGIFNα monotherapy.^9,10^ In order to improve therapeutic outcomes, combinations of IFNs and NUCs have been assessed in several clinical studies, showing that NUCs may enhance proliferating properties of HBV-specific T cells, and combined PEGIFNα and NUCs therapy could induce activation and differentiation of effector cells, and achieve higher proportion of HBsAg loss (∼9.1%) within finite duration.^11,12^ Whereas merely suppressing HBV replication by NUCs is insufficient to enhance adaptive immunity or improve IFNs response, the curative rate of CHB is still far from satisfactory, highlighting the importance of exploring novel curative modalities.

Accumulating evidences indicate that extraordinarily high levels of circulating viral antigens prevent the production of immune responses and contribute to HBV-specific immune tolerance and viral persistence.^13–15^ Thus, relieving viral antigen burden and reconstitution of antiviral function of immune system represent key factors for CHB therapy. RNA interference (RNAi), which can potently reduce viral mRNA, antigen load and viremia, has the potential to relieve HBsAg-related immunotolerance, thus representing a promising therapeutic strategy under clinical investigation.^16–19^ It has also been validated to benefit viral burden-induced cytopathology in our previous study.^20^ However, the functional cure rate is fairly low in CHB patients receiving RNAi monotreatment, since RNAi could not trigger protective antiviral immune responses directly.^21,22^ With the advantage of the AAV-HBV-based HBV carrier mice and HBV transgenic mice that mimics viral persistence and immune tolerance, recent studies have demonstrated that lowering HBV antigen load by RNAi accompanied with immune stimulation such as therapeutic vaccines is sufficient to overcome virus-specific T/B cell immune tolerance and promote HBV-specific immune reawakening.^23–25^ Additional combinational strategies based on clinical or preclinical drugs are warranted to obtain better reconstitution of antiviral immune functions and to deepen our understanding on viral persistence and immune tolerance.

Recent clinical studies showed that combining small interfering RNAs (siRNAs) with interferons can achieve deeper HBsAg decline and higher rates of HBsAg seroclearance and seroconversion, representing a promising combinational therapeutic strategy.^26–28^ However, little is known about the behavior and mechanisms of combined PEGIFNα plus RNAi therapy on the elimination of viral products and restoration of immune responses since there are lack of adequate animal models. Recently, an extracellular humanized IFNAR (IFNAR-hEC) mouse model was constructed by our group, has been validated to be responsive to human IFNs with intact endogenous immune system, and has been used to evaluate different effects of human IFN subtypes.^29^ With this model, and the in-house established GalNac-conjugated HBV siRNA (GalNac-siHBV),^30^ we were able to comprehensively investigate the effects and clarify the underlying mechanisms of combined PEGIFNα plus RNAi therapy. The results showed that combined therapy induced a potent reduction of HBsAg of ∼4log_10_ IU/mL, and achieved a HBsAg seroconversion rate of ∼30%. The synergistic efficacy of combined therapy was mainly attributed to the ablation of high viral burden-induced immune tolerance by RNAi, the induction of unspecific and virus-specific immune responses by PEGIFNα, and the altered T and B cell communications by combined therapy.

## MATERIAL AND METHODS

### Animal models

C57BL/6 mice were purchased from GemPharmatech Co., Ltd (Jiangsu, China). IFNAR-hEC mice were bled by Cyagen biotechnology company (Jiangsu, China). Mice experiments were approved by the Biomedical Research Ethics Committee of the Fudan University.

The rAAV8-HBV1.3 (ayw) viruses were purchased from Five Plus Biotech (Beijing, China). Animal models were constructed via intravenously injection with the rAAV8-HBV1.3 viruses (2.5×10^10^ viral genome (v.g.) copies per mice) and were grouped to obtain similar HBsAg levels.^31^

### Animal treatment

HBV-specific siRNAs were designed to target all forms of HBV transcripts (GalNac-siHBV). 2.5 mg/kg of GalNac-siHBV were subcutaneously (s.c.) injected every 4 weeks.

The IFNAR-hEC mice received 30 μg/kg of PEGASYS (PEGASYS; Roche) by subcutaneous injection (s.c.) twice per week for 20 weeks.

Two micrograms of HBsAg (ayw) (Fitzgerald, USA) plus 25 μg of CpG-1826 (MCE, USA) were subcutaneously injected weekly for vaccination.

### Statistical analysis

Data were analyzed using an unpaired two-tailed Student’s *t-test*, one-way analysis of variance (ANOVA), or two-way ANOVA with Sidak/Dunnett multiple comparison correction, with GraphPad Prism. Numbers in graphs indicate *P* values.

## RESULTS

### RNAi rather than PEGIFNα broke high viral antigen-induced immune tolerance

We firstly designed an efficient RNAi therapeutic agent of combined siRNA triggers targeting HBsAg (siHBs) and HBx (siHBx), respectively, to knockdown all forms of HBV transcripts with high efficiency (GalNac-siHBV) (**Fig. S1A-B**). The rAAV8-HBV1.3-based chronic HBV-carrier mouse model was utilized to evaluate the *in vivo* ^32^activity of relative agents. GalNAc-siHBV potently and dose-dependently suppressed antigenemia and viremia, whereas the positive control Entecavir (ETV) blocked HBV replication but failed to reduce antigenemia (**Fig. 1A, Fig. S1**). Neither ETV nor RNAi triggered antiviral immune responses (**Fig. S2A-D**). The clinically used PEGIFNα2a, PEGASYS, was evaluated in comparison with RNAi by utilizing IFNAR-hEC mouse model. Unlike RNAi, PEGIFNα led to moderate viral antigen reduction but evident unspecific immune activation (**Fig. 1B, Fig. S2E**). No evident reduction of HBV DNA was observed due to ∼2log_10_ lower HBV DNA levels in IFNAR-hEC mouse than in the wildtype mouse (also reported by Wang et al.).^32^ Besides, PEGIFNα treatment triggered a slight increase of HBV-specific T cell frequencies after weeks of treatment (**Fig. S2F**).

**Fig. 1.**
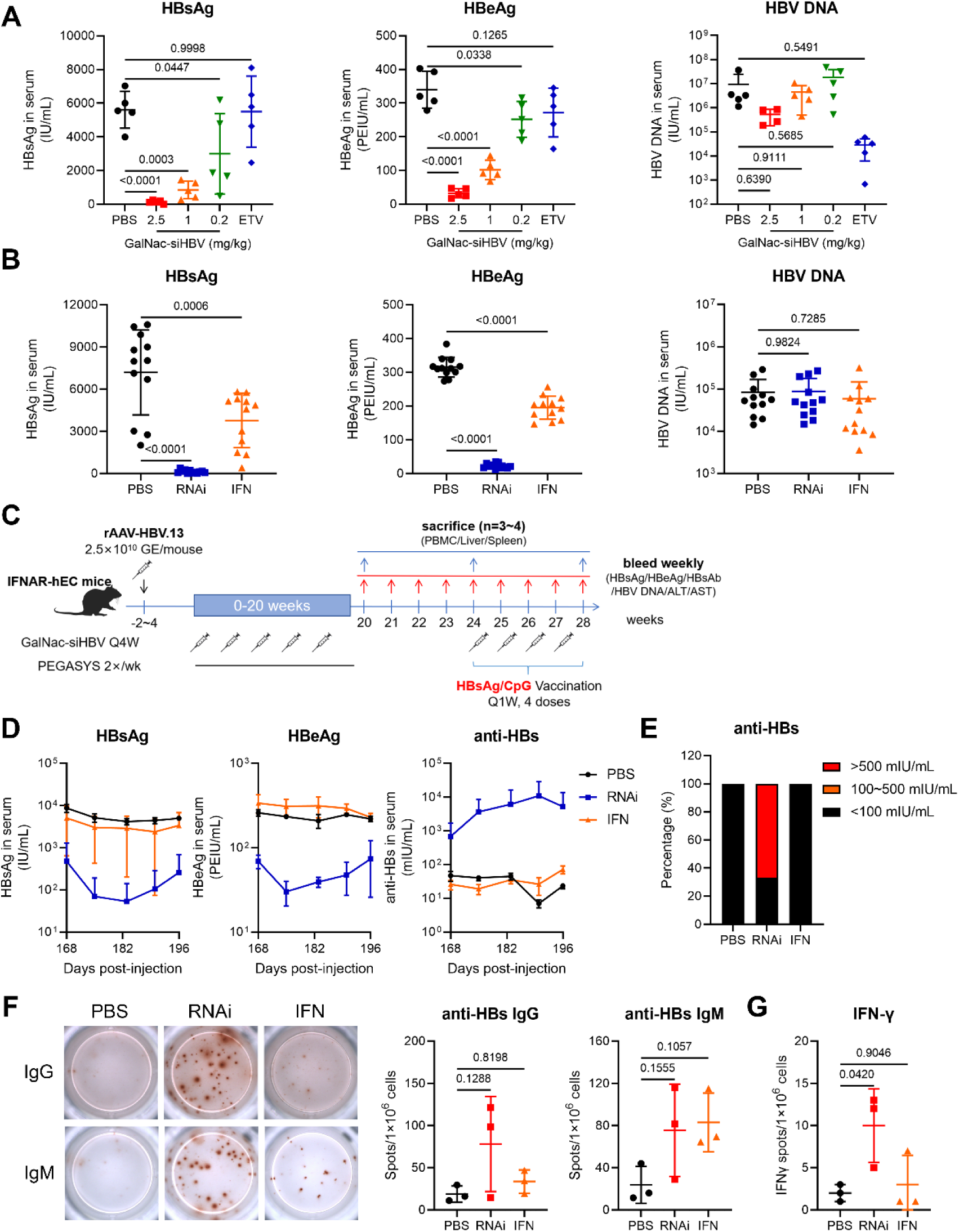
RNAi rather than PEGIFNα broke high virus antigen-related immune tolerance in chronic HBV-carrier mice. **(A)** Chronic HBV-carrier mice were treated with GalNac-siHBV (0.2/1/2.5 mg/kg, s.c., single dose), or Entecavir (1 mg/kg, per oral, daily doses). Seral HBsAg, HBeAg and HBV DNA levels were determined at day 7 (n = 5). **(B)** Chronic HBV-carrier IFNAR-hEC mice were treated with GalNac-siHBV (2.5 mg/kg, s.c.) or PEGASYS (30 μg/kg, s.c., 2×wk). Seral HBsAg, HBeAg and HBV DNA levels were determined at day 7 (n = 12). **(C)** Schematics of treatment procedures. Chronic HBV-carrier IFNAR-hEC mice were treated with GalNac-siHBV(2.5 gm/kg, s.c., Q4W) or PEGASYS (30 μg/kg, s.c., 2×wk) for 20 weeks, then treatment was stopped. Four weeks later, HBsAg/CpG-1826 vaccination was performed (Q1W, 4 doses in total). **(D)** Seral HBsAg, HBeAg and anti-HBs levels were determined (n = 3). **(E)** The percentage of HBsAg seroconversion. **(F)** Specific B cell responses to HBsAg were tested by B cell ELISpot assay at week 28. Statistical analysis were displayed (n = 3). **(G)** Liver T cell responses to HBV-specific peptides (S109 and C93) were tested by a T cell ELISpot assay (n = 3). Data were analyzed by one-way ANOVA with Dunnett multiple comparison correction (A-B, F-G), and presented as means±SD. Numbers in graphs indicate *P* values.

To questionee whether RNAi or PEGIFNα could help relieve HBV-related immune tolerance, HBsAg/CpG-1826 vaccination was applied after 20 weeks of treatment (**Fig. 1C**). Vaccination neither triggered antigen reduction nor anti-HBs antibodies secretion in PBS/PEGIFNα-treated mice, whereas RNAi treatment plus vaccination induced anti-HBs secretion in 67% (2/3) of mice (**Fig. 1D-E**). ELISpot analysis revealed enhanced anti-HBs IgG isotype secretion and increased IFN-γ secretion after *ex vivo* HBV-specific peptides stimulation in the vaccinated RNAi group (**Fig. 1F-G**). These results showed that siRNA has superiority over NUCs and PEGIFNα in reducing antigenemia and relieving high viral antigen load-related immune tolerance.

### Combined RNAi and PEGIFNα therapy exhibited synergistic antiviral effects in HBV-carrier IFNAR-hEC mice

We then inquired whether combined RNAi and PEGIFNα treatment could obtain better efficacy against HBV. Chronic HBV-carrier IFNAR-hEC mice were treated with either GalNac-siHBV (2.5 mg/kg, Q4W), PEGASYS (30 μg/kg, 2×/wk), or both drugs concurrently (**Fig. 2A**). RNAi treatment induced a significant reduction of HBsAg (- 2.18log_10_ IU/mL, vs PBS) and HBeAg at week 16. PEGIFNα alone only led to a moderate trend of HBsAg reduction (-0.51log_10_ IU/mL), with partial mice refractory to PEGIFNα treatment. Combined therapy further suppressed seral HBsAg (-4.42log_10_ IU/mL) and HBeAg levels (**Fig. 2B-D**). Similar results were observed in female mice (**Fig. S3A-D**). Specifically, combined treatment induced a higher incidence of HBsAg seroconversion (∼30% (4/12)) and higher levels of anti-HBs secretion (>1,000 mIU/mL) after 20 weeks of treatment (**Fig. 2E**).

**Fig. 2.**
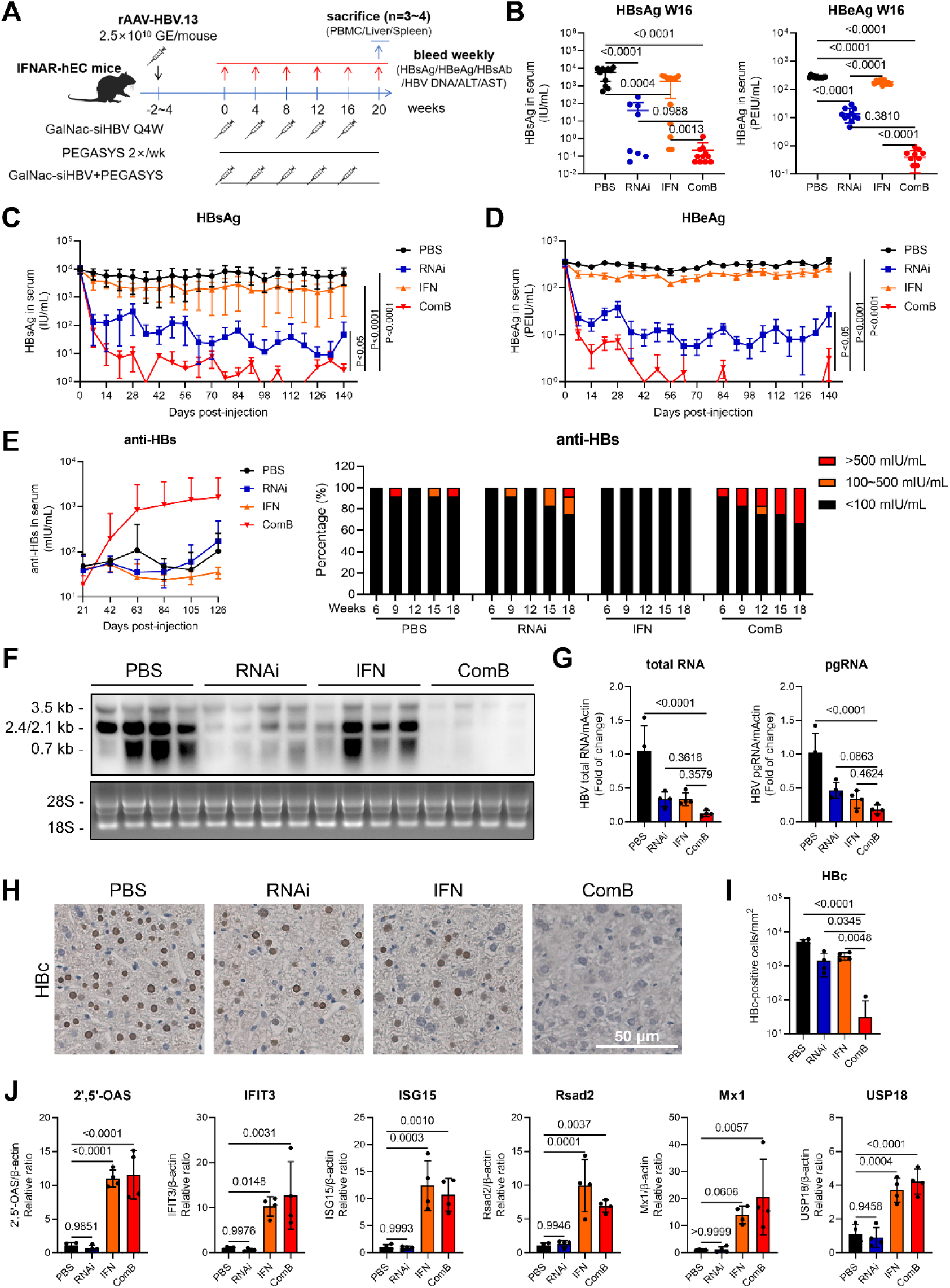
Combined RNAi and PEGIFNα treatment displayed synergistic antiviral efficacy in chronic HBV-carrier IFN-hEC mice. **(A)** Schematics of treatment procedures. Briefly, chronic HBV-carrier IFNAR-hEC mice were treated with GalNac-siHBV (2.5 mg/kg, s.c., Q4W), PEGASYS (30 μg/kg, s.c., 2×wk), or both concurrently. **(B)** Seral HBsAg and HBeAg levels at week 16 were determined. **(C)** Seral HBsAg and **(D)** HBeAg levels during 20 weeks’ treatment were evaluated. **(E)** Seral anti-HBs levels (left) and the percentage of HBsAg seroconversion (right) in mice of different groups. **(F)** Intrahepatic HBV RNA levels were determined via Norther Blot analysis and **(G)** real-time quantitative PCR (RT-qPCR) analysis (n = 4). **(H)** Representative graphs of intrahepatic HBc. Scale bar indicates 50 μm. **(I)** The HBc-positive cells per mm^2^ liver sections (n = 4). **(J)** The expression levels of interferon-stimulated genes (ISGs) relative to *actin* mRNA levels (n = 4). Data were analyzed using one-way analysis of variance (ANOVA) with Dunnett multiple comparison correction (B, G, I-J), or two-way ANOVA with Sidak multiple comparison correction (C-D), and presented as means±SD. Numbers in graphs indicate *P* values.

The intrahepatic HBV RNA and DNA load (**Fig. 2F-G, Fig. S3E-F**) and the intranuclear HBc staining (**Fig. 2H-I, Fig. S3G-H**) were remarkably diminished in combined therapy group. The successful induction of typical interferon-stimulated genes (ISGs) was confirmed via real-time quantitative PCR (RT-qPCR) analysis (**Fig. 2J, Fig. S3I**). Global transcriptional analysis of liver tissues showed that, RNAi treatment influenced metabolism-related pathways, combined therapy further enhanced immune response-related pathways (**Fig. S4**). Besides, combined therapy displayed satisfactory safety profiles, with no evident ALT/AST elevation, body weight reduction, or organism abnormality, aside from some reduction in the percentage of blood cells that attributed to PEGIFNα treatment (**Fig. S5**). Together, these results showed that combined RNAi and PEGIFNα therapy could be applied as an effective treatment to exhibit synergistic antiviral effects along with high security profiles.

### Combined RNAi and PEGIFNα therapy led to augmented HBV-specific T cell responses

The peripheral blood mononuclear cells (PBMCs) were investigated by flow cytometry (FACS) analysis during the 20 weeks’ treatment. PEGIFNα mono- and combined-treatment resulted in significantly increased expression of Ki67, CD69 and CD38 markers on CD8^+^ and CD4^+^ T cells, the effects of which were more pronounced on CD8^+^ than CD4^+^ T cells (**Fig. 3A-B**). The expression levels of Ki67 and CD69 molecule on NKs/DCs were also increased by PEGIFNα treatment (**Fig. S6B-D**). Whereas the magnitude of T/NK/DC cell activation by PEGIFNα did not correlate with seral HBsAg levels, this was due to the absence of a significant difference between the PEGIFNα only and the combined groups in terms of immune activation. In addition, an increased proportion of Foxp3^+^ CD25^+^ regulatory T (Treg) cells in the circulation were induced by PEGIFNα treatment (**Fig. S6A**).

**Fig. 3.**
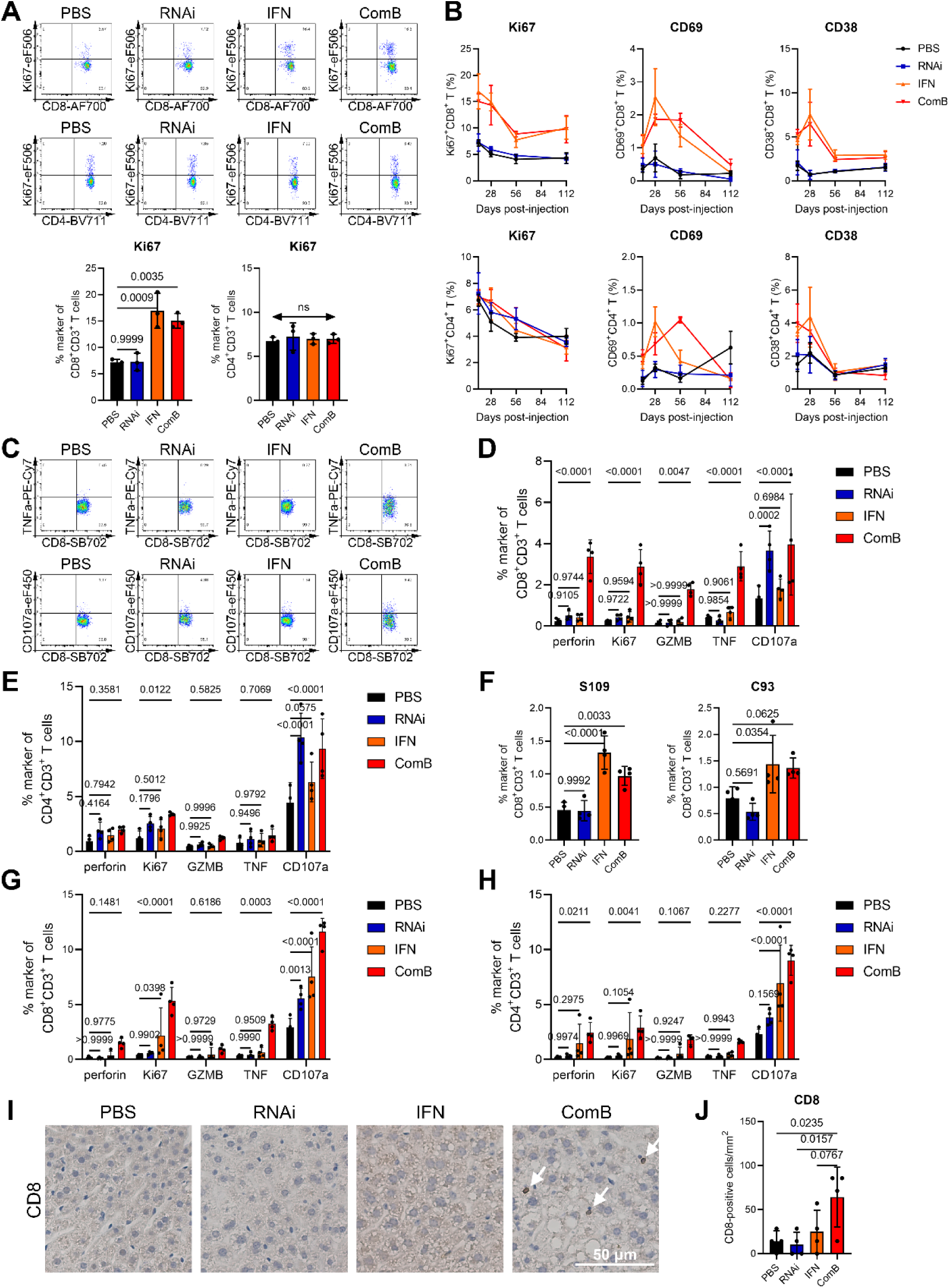
Combined RNAi and PEGIFNα treatment augmented HBV-specific CD8^+^ and CD4^+^ T cell responses. **(A)** The proportion of Ki67-positive CD8^+^CD3^+^ and CD4^+^CD3^+^ T cells in PBMCs (n = 3). **(B)** The percentage of Ki67, CD69 and CD38-positive CD8^+^ and CD4^+^ T cells at indicated time points (n = 3). **(C)** Representative density plots and **(D)** statistical analysis of the proportion of cells in intrahepatic CD8^+^CD3^+^ T cells and **(E)** CD4^+^CD3^+^ T cells (n = 4). **(F)** The proportion of HBsAg-and core-specific CD8^+^ T cells in gated liver lymphocytes (n = 4). **(G)** Statistical analysis of the proportion of perforin, Ki67, GZMB, TNFα, IFNγ and CD107a expressing splenic CD8^+^CD3^+^ and **(H)** CD4^+^CD3^+^ T cells after *ex vivo* HBV-derived peptide stimulation was displayed (n = 4). **(I)** Immunohistochemical analysis of intrahepatic CD8-positve cells. Scale bar indicates 50 μm. **(J)** Statistical analysis of the CD8-positve cells per mm^2^ liver sections (n = 4). Data were analyzed by one-way ANOVA with Dunnett multiple comparison correction (A, F, J), or two-way ANOVA with Sidak multiple comparison correction (D-E, G-H), and were presented as means±SD. Numbers in graphs indicate *P* values.

The peripheral T cells of combined therapy exhibited higher functionality, with higher percentage of HBsAg- and core-specific T cells, as well as more CD8^+^ and CD4^+^ T cells stained positive with CD107a, TNF-α, granzyme B (GZMB) and interferon-γ (IFN-γ) (vs PBS) (**Fig. S6E-G**). Higher percentages of CD8^+^ and CD4^+^ T cells were positively stained with perforin, Ki67, GZMB, TNF-α and CD107a upon *ex vivo* HBV-derived peptides restimulation in intrahepatic lymphocytes of combined therapy, and higher proportions of HBsAg- and core-specific CD8^+^ T cells were detected (vs PBS) (**Fig. 3C-F**). Similar results were observed in splenocytes, showing enhanced CD8^+^ and CD4^+^ T cell functionality in mice of combined therapy (**Fig. 3G-H**). Immunohistochemical (IHC) analysis showed an evident intrahepatic infiltration of CD8^+^ T cells in mice of combined therapy (**Fig. 3I-J**). Together, these results showed that mice of combined RNAi and PEGIFNα treatment gained enhanced unspecific and HBV-specific T cell functionality, which could be important for the synergistic mechanisms in antagonizing chronic HBV infection.

### Combined RNAi and PEGIFNα therapy improved HBV-specific B cell immune responses

Analysis of B cell phenotypes revealed that combined RNAi and PEGIFNα treatment induced a higher percentage of CD19^-^CD138^+^ plasma B cells, CD27^+^CD38^-^ memory B cells, as well as higher percentage of IgD^-^IgM^-^ class switched B cells, which is closely correlated with antibody secretion (**Fig. 4A-B**). Combined therapy also induced higher expression levels of Ki67, MHC-II, CD86 and CD80 activation molecule on CD19^+^CD3^-^ B cells (vs PBS) (**Fig. 4C-E**). However, the percentage of HBsAg-specific B cells did not change (**Fig. 4B**). The phenotypes and functionality of HBsAg-specific B cells in different groups were evaluated, showing that mean expression levels of CD80, Ki67 and CD11c molecule were significantly elevated on HBsAg-specific B cells of combined therapy, mirroring the trend of global B cells (**Fig. 4F**). These results together confirmed that combined RNAi and PEGIFNα treatment improved the functionality of global and HBsAg-specific B cells, thus promoting HBsAg seroconversion in treated mice.

**Fig. 4.**
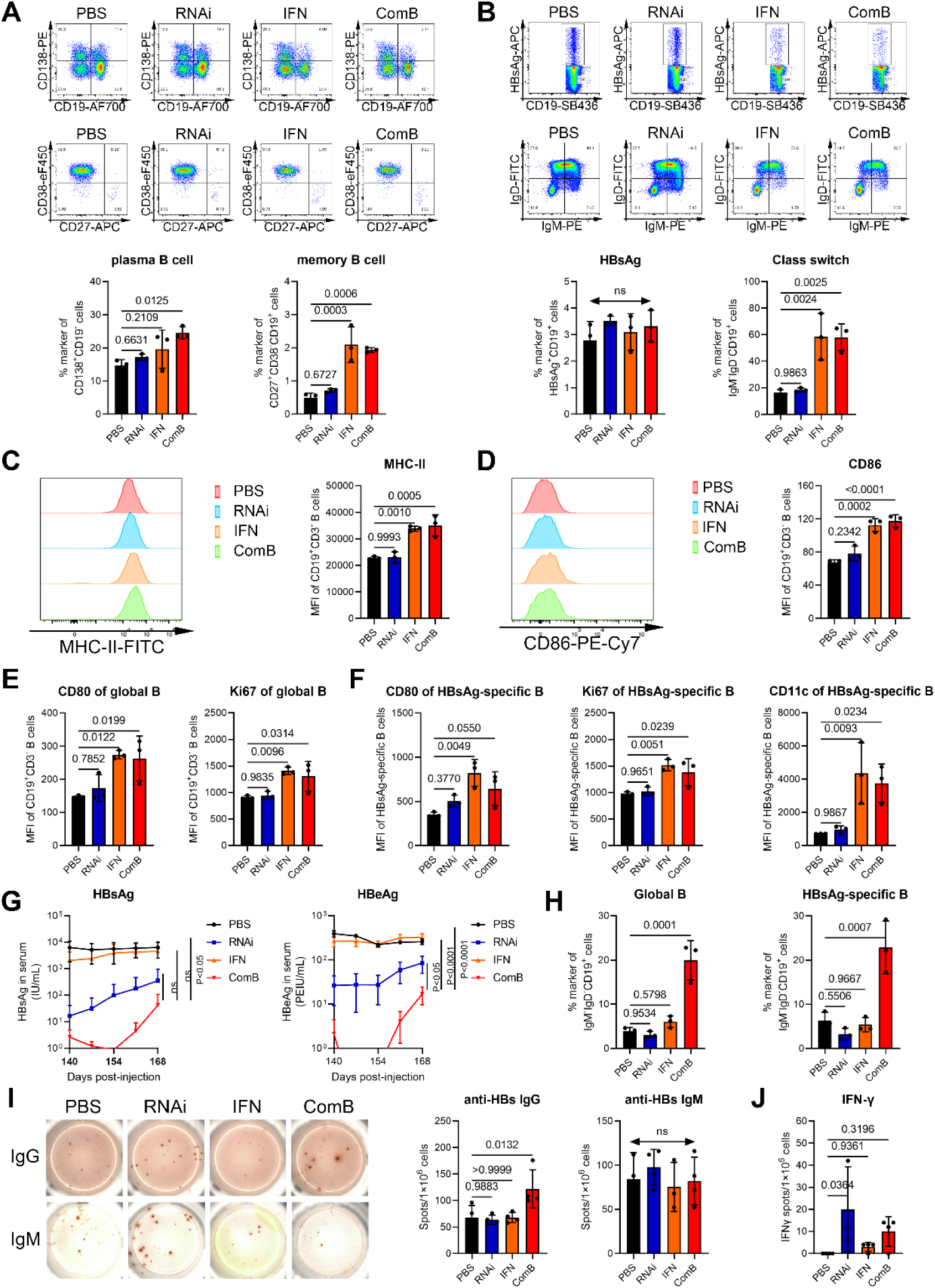
Combined RNAi and PEGIFNα treatment triggered prolonged B cell activation and class switch. **(A)** The percentage of CD19^-^CD138^+^ plasma B cells and CD27^+^CD38^-^ memory B cells (n = 3). **(B)** The proportion of HBsAg-specific and IgD^-^IgM^-^ classed switched B cells (n = 3). **(C)** Mean fluorescent intensity of MHC-II and **(D)** CD86 staining in gated CD19^+^CD3^-^ B (global B) cells (n = 3). **(E)** Statistical analysis of mean fluorescent intensity (MFI) of CD80 and Ki67 staining in global B cells (n = 3). **(F)** CD80, Ki67 and CD11c MFI in HBsAg-specific B cells (n = 3). **(G)** Mice were treated with GalNac-siHBV with or without PEGASYS co-treatment for 20 weeks, then treatment was stopped. Seral HBsAg and HBeAg levels. **(H)** The proportion of the IgD^-^IgM^-^ classed switched B cells in global B (left) and HBsAg-specific B cells (right) (n = 3). **(I)** Specific B cell responses to HBsAg were tested in a B cell ELISpot assay at week 28 (n = 4). **(J)** Liver T cell responses to HBV-specific peptides (S109 and C93) were tested by a T cell ELISpot assay (n = 4). Data were analyzed by one-way ANOVA with Dunnett multiple comparison correction (A-F, H-J), or two-way ANOVA with Sidak multiple comparison correction (G), and were presented as means±SD. Numbers in graphs indicate *P* values.

We further inquired whether combined treatment could obtain prolonged viral control and immune response, treatment was continued for 20 weeks then stopped for 4 weeks. HBV antigens demonstrated a trend of rebound, but the inhibition rates of HBsAg retained over 2log_10_ IU/mL (vs PBS) in mice of combined therapy (**Fig. 4G**). Specifically, the percentage of IgD^-^IgM^-^ class switched B cells retained at higher level in combined therapy groups at week 24, both in global and HBsAg-specific B cells (**Fig. 4H**). Combined treatment also induced an evident elevation of HBsAg-specific IgG but not IgM antibody secretion in splenocytes (**Fig. 4I**). Liver lymphocytes of combined therapy secreted higher levels of IFN-γ when stimulated with HBV-specific peptides *ex vivo* (**Fig. 3J**). RNA-seq analysis of splenic B cells at week 20 and week 24 showed that, PEGIFNα treatment exhibited potent effects on inducing B cell activation (**Fig. S7**), with the combined therapy induced prolonged B cell responses after treatment cessation (**Fig. S8**). Together, these results showed that the antiviral effects and immune regulatory effects of combined therapy could maintain for at least 1 month after treatment stopped.

### Single-cell RNA sequencing revealed distinct destiny of T and B cells after long-term RNAi and PEGIFNα therapy

Comprehensive unbiased scRNA-seq analyze of intrahepatic Cd45^+^ immune cells at week 20 demonstrated high-quality expression data for 44,603 cells after quality control (**Fig. 5A**). Six different cell clusters were assigned based on differential marker gene expression, including hepatocytes, monocytes, neutrophils, T cells, B cells and NK cells (**Fig. 5B, Fig. S9**). An evident change in the proportion of immune cells across different conditions was observed. PEGIFNα treatment induced an increased proportion of T cells, NKs, and neutrophils, and a decreased proportion of B cells (**Fig. 5C-D**). FACS analysis confirmed increased T but reduced B cell proportions (**Fig. 5E**). Violin plot analysis showed that PEGIFNα treatment induced an evident upregulation of apoptosis genes like Bax, Cycs, S100a8 and S100a9 (**Fig. 5F**). Reduced B cells and improved neutrophils infiltration in intrahepatic environment by PEGIFNα treatment were confirmed by reanalyzing scRNA-seq data of huIFNAR mice (GSE237519) (**Fig. S13**). Further sub-clustering and annotation with known marker genes revealed the complexity of T and B cells, with much more Cd8^+^ T sub-cluster cells than Cd4^+^ T cells, including naïve-like (Cd8-C1, Cd8-C3, Cd8-C4, Cd8-C5), cytotoxic (Cd8-C2), proliferating (Cd8-C6) and activated Cd8^+^ T cells (Cd8-C7) (**Fig. 6G-H, Fig. S10A, C**). The B cells were sub-clustered into naïve (B-C1, B-C2, B-C3, B-C9), transitional (B-C4), atypical memory (atMBC) (B-C5), marginal zoom (MZB) (B-C6), classical memory (MBC) (B-C7), and activated B cells (B-C8) (**Fig. 6I-J, Fig. S10B, D**).

**Fig. 5.**
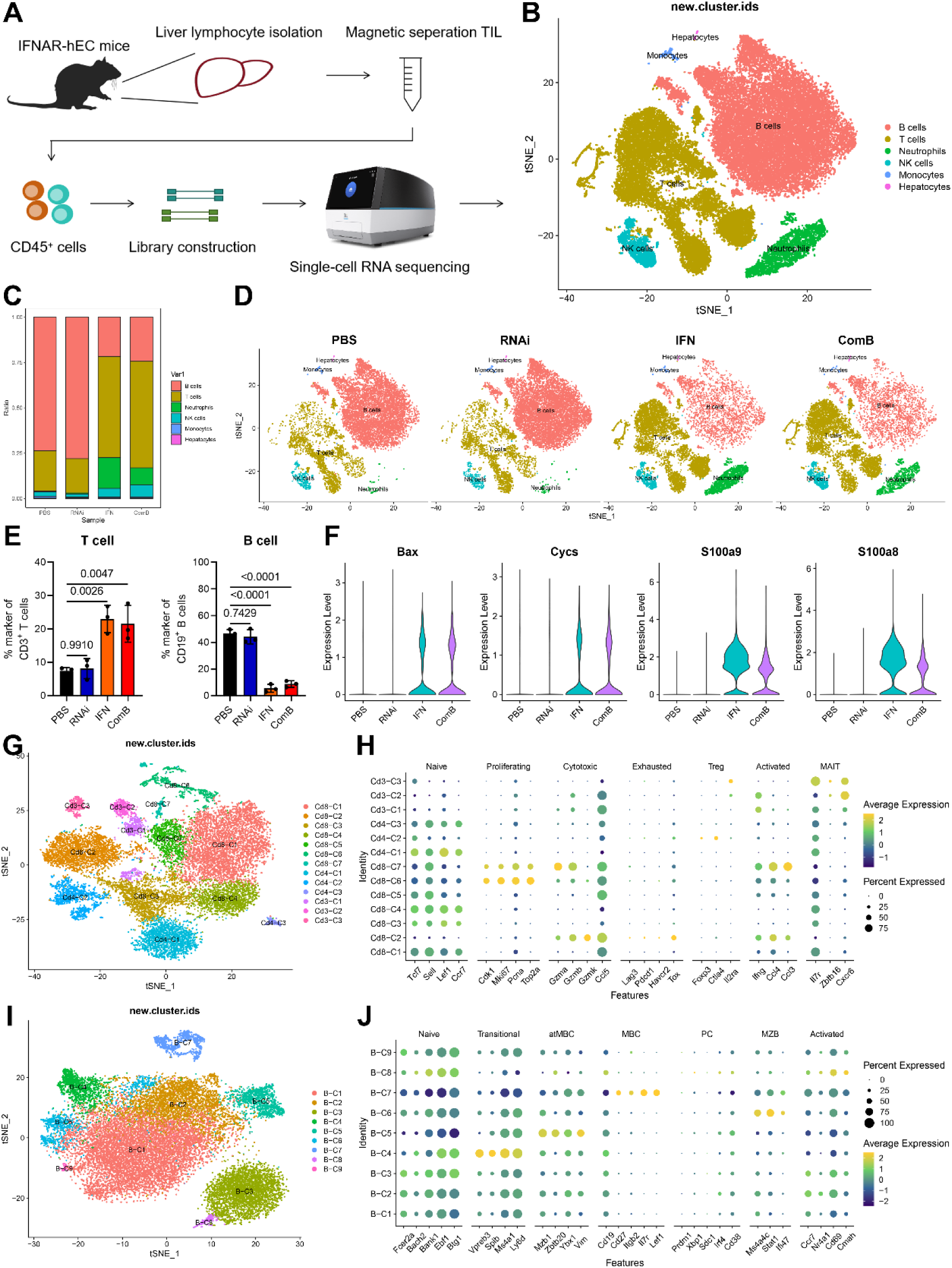
Single-cell RNA sequencing revealed the complexity of T and B cells in combined RNAi and PEGIFNα treated mice. **(A)** Schematics of experiment pipeline. Mice intrahepatic Cd45^+^ immune cells were collected for scRNA-seq at week 20. **(B)** TSNE plot of intrahepatic immune cells colored according to cell types. **(C)** The proportion of each cluster in different groups. **(D)** TSNE plot of intrahepatic immune cells from mice of different treatment. **(E)** The percentage of CD3^+^CD19^-^ T cell and CD19^+^CD3^-^ B cells in gated lymphocytes at week 20 (n = 3). **(F)** Violin plots showing the expression levels of Bax, Cycs, S100a9 and S100a8. **(G)** TSNE plot of identified T cell subsets. **(H)** Dot plots showing the expression of marker genes in T cell subsets. **(I)** TSNE plot of identified B cell subsets. **(J)** The expression of marker genes in B cell subsets. Data were analyzed by one-way ANOVA with Dunnett multiple comparison correction (E), and were presented as means±SD. Numbers in graphs indicate *P* values.

**Fig. 6.**
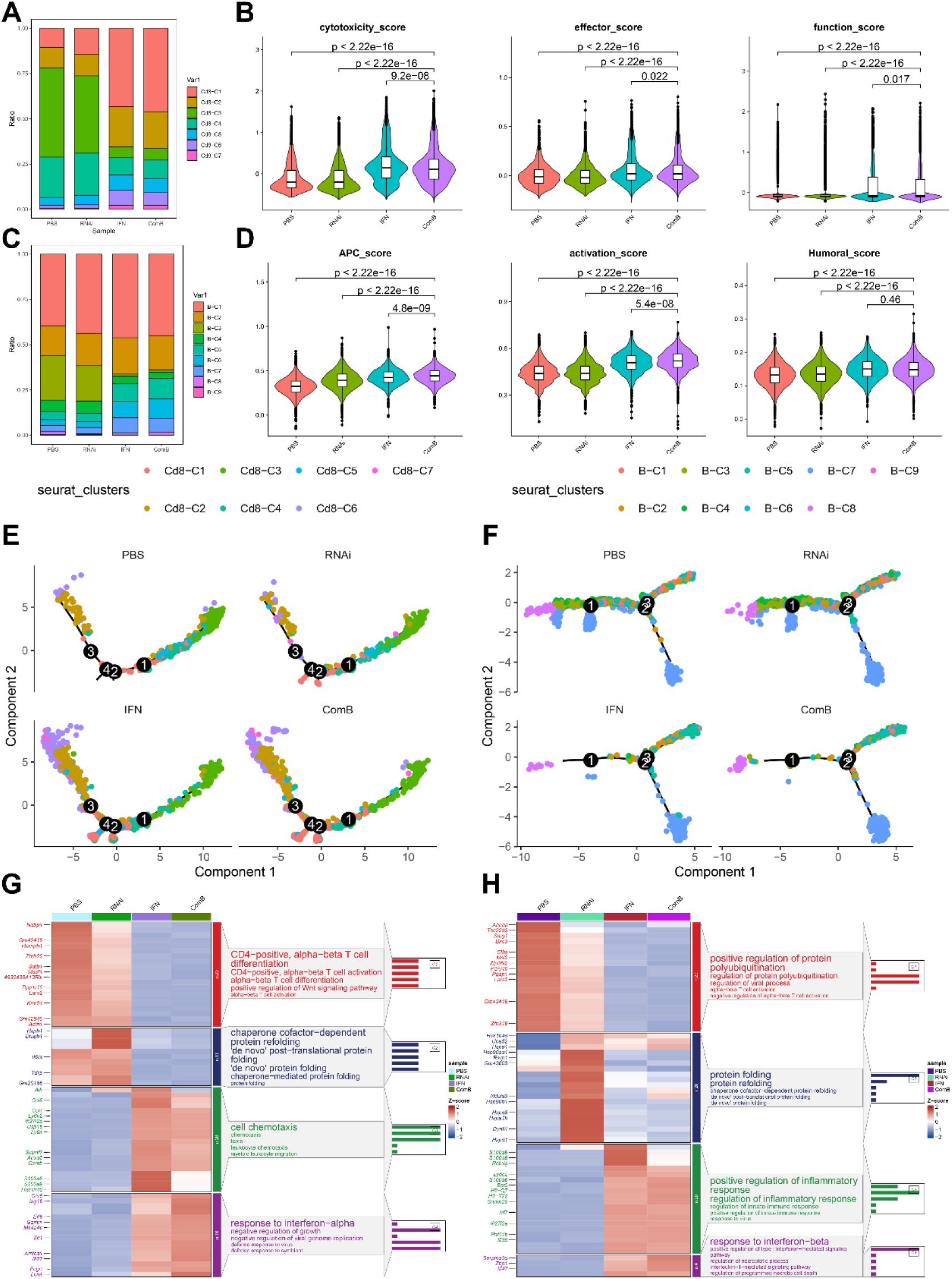
Assessment of the functional states of liver lymphocytes in combined RNAi and PEGIFNα treated mice. **(A)** The proportion and **(B)** violin plots of gene set enrichment analysis scores of CD8^+^ T cell subclusters, the cytotoxicity, effector and function-related scores were displayed. **(C)** The proportion and **(D)** violin plots of gene set enrichment analysis scores of B cell subclusters, the antigen processing and presenting, activation and humoral immune response-related scores were calculated. **(E)** Pseudo-time trajectory analysis of CD8^+^ T and **(F)** B cells showing the distribution of each cell subset on the development trajectory. **(G)** Heatmap and pathway enrichment analysis of different expressed genes in CD8^+^ T and **(H)** B cells by ClusterGVis. Data were analyzed by unpaired two-tailed Students’ *t*-test analysis (B, D), and were presented as means±SD. Numbers in graphs indicate *P* values.

Further analysis revealed that combined treatment led to an elevation of the percentage of Cd8-C1, Cd8-C2, Cd8-C5, Cd8-C6 and Cd8-C7 cells, as well as induced a significant elevation of cytotoxic, effector and function-related gene scores in Cd8^+^ T cells (**Fig. 6A-B**). The proportion of B-C3 subcluster reduced with the proportion of B-C5, B-C6 and B-C7 subclusters increased in combined treatment groups, with the antigen processing and presentation, activation, and humoral immune response-related gene scores of B cells were elevated (**Fig. 6C-D**). This phenomenon was also observed in huIFNAR mice (**Fig. S14A-B**), and was consistent with that observed in clinical patients.^28^ The pseudo-time analysis confirmed the enhanced development of Cd8^+^ T cells from naïve-like to proliferating and cytotoxic phenotypes by combined therapy (**Fig. 6E, Fig. S11A-B**). There was also an enhanced development of B cells from naïve-like to transitional or memory-like B cells by combined therapy (**Fig. 6F, Fig. S11C-D**). Pathway enrichment analysis revealed that, RNAi treatment improved signaling pathways associated with protein folding, PEGIFNα induced inflammatory response and cell chemotaxis related pathways, whereas combined treatment further enhanced interferon-α/β responses and antiviral responses in T and B cells (**Fig. 7G-H, Fig. S12A**).

**Fig. 7.**
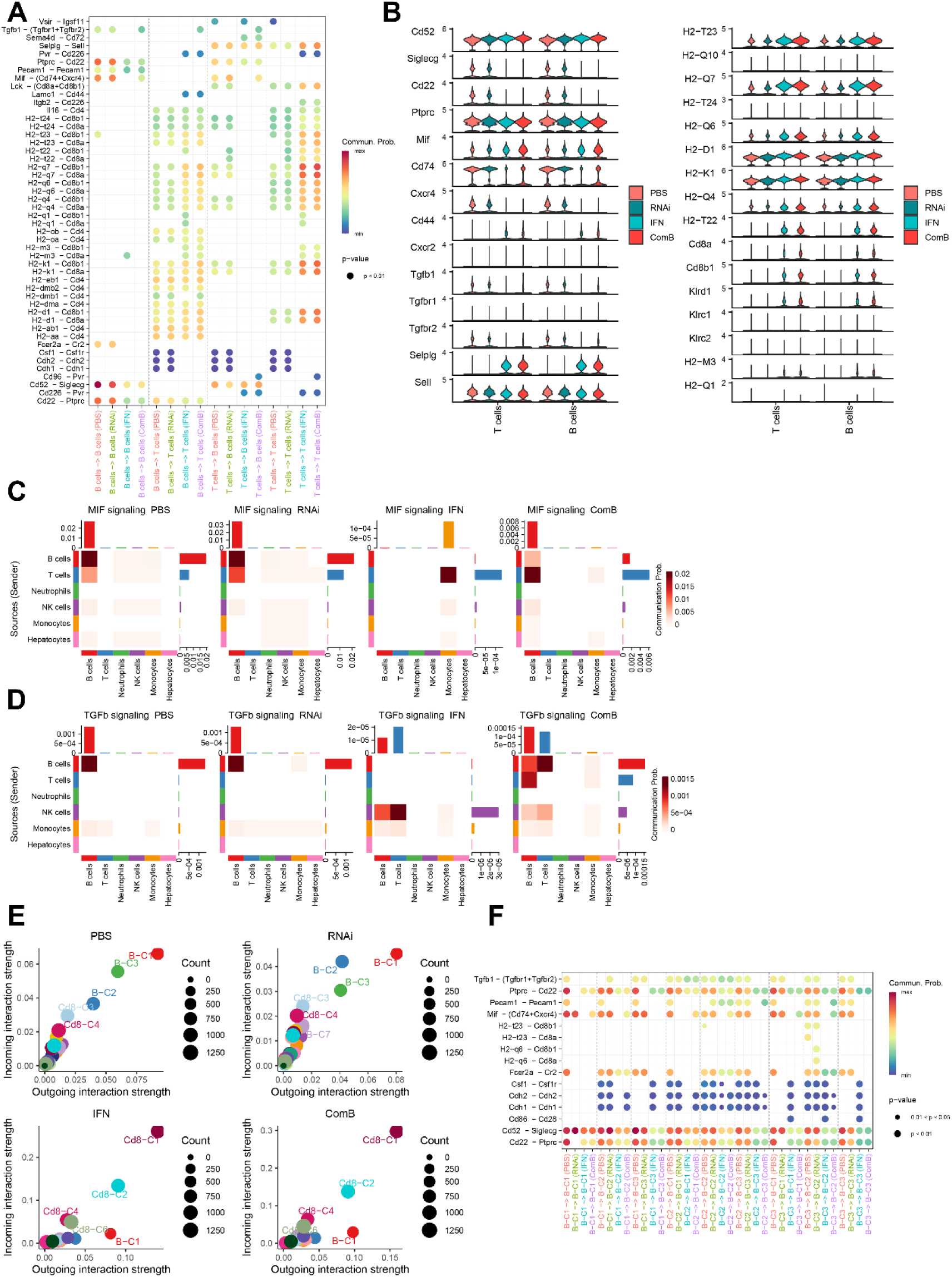
Cell-cell communication analysis indicated the complex interactions between T and B cells after combined RNAi and PEGIFNα treatment. **(A)** Comparison of the significant ligand-receptor pairs between T and B cells. Dot color represents the communication probability of the specific ligand-receptor pair between sender and receiver clusters. **(B)** Violin plots showing the expression levels of ligand-receptor pairs associate with T cells and B cells crosstalk in different groups. **(C)** Heatmap showing communication probability of MIF and **(D)** TGFb signaling. The top and right bar around the heatmap represent the sum of column values (incoming signaling) and row values (outgoing signaling) respectively. **(E)** Incoming and outcoming strengths of the cell interaction signals of T and B cell subpopulations. **(F)** Comparison of the significant ligand-receptor pairs across different groups among B cell subclusters.

### Combined RNAi and PEGIFNα therapy changed the interaction modes between T and B cells

Communication analysis of individual samples showed a dominant crosstalk between T and B cells, via H2-aa/ab1 ligand and CD4 receptor pairs (MHC-II signaling), promoting antigen processing and presentation, and PEGIFNα treatment further enhanced T cells and T cells interactions mainly through H2-q7 ligand and CD8a/b1 receptor pairs (MHC-I signaling) (**Fig. S12B-C**). Communications across different groups were further evaluated. Cd22-Ptprc and Cd52-Siglecg pairs, which function as negative B cell signaling modulators, mainly medicate B cell-B cell communication, indicating an inhibitory immune state of B cells.^33,34^ Specifically, PEGIFNα treatment reduced Cd22-Ptprc and Cd52-Siglecg pair interactions by depressing ligand or receptor expression (**Fig. 7A-B**). Combined treatment also enhanced Tgfb1-(Tgfbr1+Tgfbr2) and Mif-(Cd74+Cxcr4) pair interactions between B cell-B cell and B cell-T cell by promoting ligand or receptor expression. Combined treatment further enhanced the MHC-I signaling between T and T cells, and improved the expression levels of MHC-I-related molecules, like H2-T23, H2-Q7, H2-Q6, H2-D1, H2-K1, H2-Q4, H2-T22, and Cd8a/b1, promoting antigen presentation (**Fig. 7B**). The MIF signaling, which promotes B cell migration but functions as a macrophage migration inhibitory factor, were changed from B cell-B cell interaction to T cell-monocyte communication by PEGIFNα treatment, but were enhanced between T and B cells by combined therapy, promoting B cell migration (**Fig. 7C**). The TGFb signaling, which mediates cell proliferation, was changed from B cell-B cell to NK-T cell communications by PEGIFNα treatment, and were increased between B and T cells by combined treatment (**Fig. 7D**). These results indicates that combined therapy may improve HBsAg antigen recognition and antibody secretion by enhancing B cell migration and T cell proliferation.

The interactions between T and B cell subclusters were analyzed, showing that the incoming and outgoing interaction strength of the B-C1, B-C2 and B-C3 cells were the most evident in PBS/RNAi treated groups, PEGIFNα treatment reduced the interaction strength among B cells but improved the interaction signaling in Cd8-C1 and Cd8-C2 T cells (**Fig. 7E**). Cell-cell communication probabilities further confirmed reduced inhibitory Cd22-Ptprc and Cd52-Siglecg pair interactions among B cells by combined therapy, leading to B cell activation (**Fig. 7F**). In addition, the Mif-(Cd74+Cxcr4) pair interaction was enhanced in combined treatment group, compared with PEGIFNα mono-treatment. Reduced inhibitory interactions among B cells and enhanced MHC-I signaling among Cd8^+^ T cells by PEGIFNα were also observed in huIFNAR mice (**Fig. S14C-E**). Together, these results showed that combined therapy reduced the inhibitory interactions between B cells and B cells, enhanced the MHC-I signaling among T cells, and improved T cell-B cell crosstalk, promoting B cell migration and T cell proliferation, thus promoting antigen clearance and immune control of the virus.

### Enhanced MHC-II-signaling networks promoted HBsAg seroconversion after combined RNAi and PEGIFNα treatment

Serum markers of typical mice in different groups were displayed. HBV antigen rebounded evidently in HBsAg-untransformed mice, but retained at undetectable levels one month after treatment cessation in HBsAg-transformed mice (**Fig. 8A**). Persistent control of HBeAg were not observed, indicating that the persistent HBsAg loss were attributed to immune control, but not cccDNA silence or clearance. We further evaluated the intrahepatic Cd45^+^ immune phenotypes after 20 weeks of combined treatment in untransformed (ComB1) and transformed (ComB2) mice (**Fig. 8B, Fig. S15A-B**). Mice of HBsAb^+^ displayed a marked reduction of infiltrated neutrophils. Further sub-clustering of T and B cells revealed an increased percentage of naïve like Cd8^+^ T and B cells, but a reduced percentage of cytotoxic, proliferating Cd8 T cells, and atMBCs (**Fig. S15C-H**). The expression levels of MHC-I-related molecules were comparable, whereas the expression levels of MHC-II-related molecules were increased, including H2-Aa, H2-Ab1, H2-Eb1, H2-DMb2, H2-Oa and H2-Ob, in HBsAg-transformed mice (**Fig. 8D**). The MHC-II signaling network between B cells and hepatocytes, and the MIF signaling between T and B cells were improved in mice of HBsAb^+^ (**Fig. 8E**). This phenomenon was consistent with that observed in clinical samples, showing an enhanced MHC-II-expressing hepatocytes in patients of functional cure.^35^

**Fig. 8.**
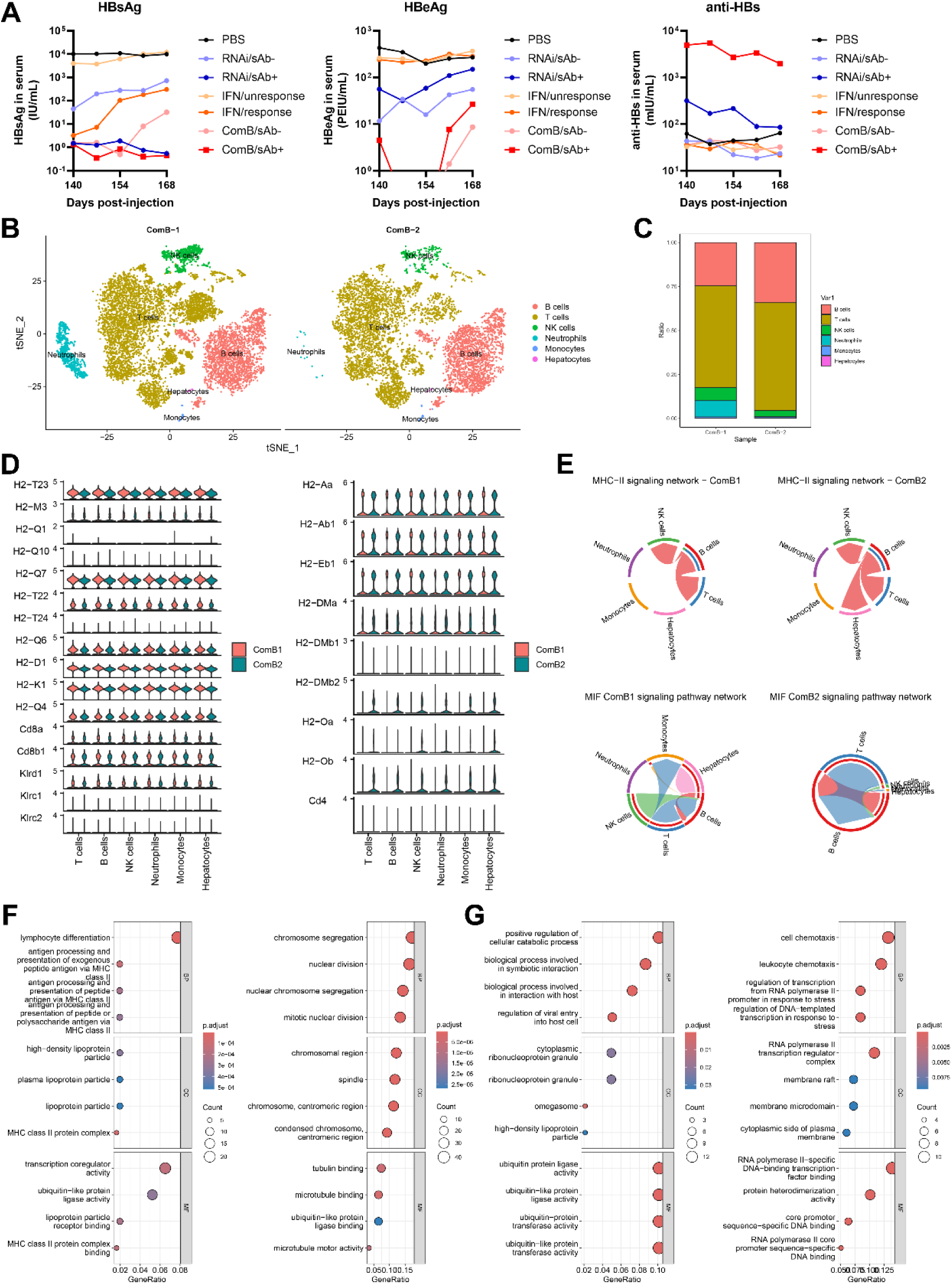
Enhanced MHC-II-signaling networks mediated HBsAg seroconversion after combined RNAi and PEGIFNα treatment. **(A)** Serum markers of typical mice in different groups (one line represented one mice). **(B)** TSNE plot of intrahepatic Cd45^+^ immune cells, colored according to cell types (ComB-1: HBsAg-untransformed; ComB-2: HBsAg-transformed). **(C)** The proportion of each cluster in different groups. **(D)** Violin plots showing the expression levels of ligand-receptor pairs associate with MHC-I (left) and MHC-II (right) signaling. **(E)** Interaction networks emphasize cell-to-cell interactions via MHC-II (top) and MIF (bottom) pathways. **(F)** Up (left) and down (right) regulated genes in Cd8^+^ T and **(G)** B cells between HBsAg-transformed and untransformed mice were subjected to pathway enrichment analysis using Gene Ontology (GO) function analysis (BP: biological process; CC: cellular component; MF: molecular function).

Pathway enrichment analysis showed that, the Cd8^+^ T cells of transformed mice displayed enhanced signaling pathway in “lymphocyte differentiation”, and “antigen processing and presentation of peptide via MHC-II”, and reduced “chromosome segregation” and “nucleic division” signaling (**Fig. 8F**). Further analysis of B cells showed an increased signaling in “positive regulation of cellular catabolic process”, and “ubiquitin protein ligase activity”, representing an improved B cell metabolic activity, and a depressed signaling in “cell chemotaxis” and “signaling related to RNA polymerase II-specific transcription”, showing a reduced trend of B cell migration and transcription. Together, these results suggested that the enhanced MHC-II signaling network between B cells and hepatocytes/Cd8^+^ T cells, and the enhanced antigen processing and presentation pathways in Cd8^+^ T cells promoted HBsAg seroconversion after long-term RNAi plus PEGIFNα treatment.

## DISCUSSION

Potent nucleos(t)ide analogues efficiently suppress viral replication but do not target HBV transcripts and cccDNA. IFN-α has the benefit of finite treatment duration but the functional cure rate is still low. HBV siRNAs represent a promising therapeutic approach currently under investigation since it can both reduce antigen and viraemia levels.^36,37^ However, merely RNAi cannot reconstruct antiviral immune responses. The viral antigens reappeared once upon treatment withdrawal. Recent studies showed that reducing HBV antigens via RNAi followed by immune stimulation with therapeutic vaccines could establish effective antiviral immune responses.^38^ In the present study, we found that siRNA has superiority over NUCs and PEGIFNα in reducing viral antigens burden and high viral antigen load-related immune tolerance, as evidenced by induced anti-HBs IgG antibodies and increased *ex vivo* peptide-stimulated IFN-γ secretion in vaccinated RNAi treatment mice, thus offering the opportunity of immune reawakening.

PEGIFNα has been demonstrated to have diverse effects on innate and adaptive immune cells during infection, and has potential to induce HBsAg seroconversion in partial CHB patients in clinical practice.^39^ It is believed to allow restoration of HBV-specific T cell function in CHB through inhibiting viremia and antigenemia and by exerting direct immunomodulating effects, whereas to what extent IFN-α can restore T cell function remains open questions.^9^ Previous studies identified that IFN-α can act directly on both CD4^+^ and CD8^+^ T cells, increasing ratios of long-lived naïve/memory T cells and enhancing effector T cell cytotoxicity.^40^ However, the immunomodulatory effects of IFN-α might be partly blunted by its antiproliferative effects and the inherent immune exhaustion in CHB patients. Based on our recently constructed IFNAR-hEC mouse model, we found that IFN-α can induce significant activation and proliferation of T cells, the effects of which were more evident in CD8^+^ than CD4^+^ T cells. Activated NKs and DCs showed upregulated CD69 and Ki67 molecules. The role of T cells in achieving HBV cure has been well-investigated, while little is known about the role of B cells in IFN-α-mediated anti-HBV immune responses. Recent studies indicated that type I IFNs can promote B cell activation and class switch during viral infection.^41^ IFNs can also facilitate antigen presentation to B cells, resulting in stronger antigen-specific responses.^42^ However, IFNs may also impair the survival and development of precursor and immature B cells, block B cell responses, and lead to the production of immunosuppressive molecules.^43^ Consistently, our results also showed that PEGIFNα treatment improved the function of B cells by upregulating MHC-II, CD80, CD86 molecules, promoting B cell proliferation, and induced high percentage of CD19^-^ CD138^+^ plasma B cells, CD27^+^CD38^-^ memory B cells, and IgD^-^IgM^-^ class switched B cells. Whereas PEGIFNα monotreatment failed to trigger anti-HBs antibody production even after 20 weeks’ treatment. Subsequent HBsAg/CpG-1826 vaccination did not trigger HBsAg-specific antibody responses and HBV-specific T cell responses as well. The percentage of FoxP3^+^CD25^+^ Treg cells was also elevated, indicating an immunoinhibitory effects of PEGIFNα.

Combined RNAi and PEGIFNα therapy have been under clinical study recently. Participants of CHB were treated with VIR-2218 (an investigational HBV siRNA) with or without PEGIFNα co-treatment. All participants of combined therapy achieved deeper suppression of viral parameters, higher rate of HBsAg seroclearance (∼14.9%, 11/69), and higher incidence of HBsAg seroconversion (∼14.5%, 10/69).^26,27^ However, the underlying synergistic mechanisms remain obscure. Using the IFNAR-hEC mice, we found that the GalNac-siHBV showed potent and dose-dependent anti-HBV activity, PEGIFNα only led to moderate suppression of viral antigens. Combined RNAi and PEGIFNα treatment induced deeper reduction of HBsAg (∼4log_10_ IU/mL, vs PBS), and induced a higher rate of anti-HBs secretion in ∼30% (4/12) of treated mice, along with higher anti-HBs titers of >1,000 mIU/mL. Mechanistically, combined therapy provided a positive synergistic response by inducing immune activation, as evidenced by improved HBV-specific T cell function after *ex vivo* peptides stimulation, and prolonged B cell class switch and anti-HBs IgG isotype secretion. Specifically, the frequency of circulating HBsAg-specific B cells did not change, but their functionality is influenced by HBsAg levels and by PEGIFNα treatment, the results of which were consistent with previous reports.^44^

scRNA-seq analysis showed that PEGIFNα treatment boosted high levels of CD8^+^ T cell responses, leading to the development of CD8^+^ T from naïve-like to proliferating and cytotoxic T cells, and enhanced communications among CD8^+^ T cells through MHC-I signaling. Whereas the percentage of naive B cells decreased with the proportions of transitional, atypical and classical memory B cells increased. Besides, the antigen processing and presentation, activation, and humoral immune response-related scores of B cells were improved. The inhibitory Cd22-Ptprc and the Cd52-Siglecg pair interactions among B cells were reduced by PEGIFNα treatment through inhibiting relative ligand or receptor expression. Combined treatment enhanced the MIF and TGFb signaling between T and B cells, promoting B cell migration and T cell proliferation. Enhanced MHC-II signaling networks between B cells and hepatocytes/Cd8^+^ T cells further promoted HBsAg seroconversion in combined treatment group. Together, these results confirmed that combined RNAi and PEGIFNα treatment exerted synergistic antiviral efficacy and orchestrated protective cellular immune effects in T cell- and B cell-dependent manners.

Although this study has some important implications for futural clinical CHB therapy, there are som<u>e</u> limitations that need to be addressed in further studies. Since the rAAV8-HBV1.3-based mouse model is not driven by the cccDNA reservoir and cannot fully reflect human immune features, the direct effects of combined treatment on cccDNA were elucidated in another study based on our recombinant cccDNA models.^45^ Considering the presence of integrated HBV DNA as a source of HBsAg production, and mice models could not fully recapitulate the phenomenon in human HBV infection, an in-depth analysis of anti-HBV and immunomodulatory mechanisms of combined therapy, based on clinical-derived samples or human liver-chimeric mice, should provide more detailed information. Besides, the seroconversion rate of HBsAg in combined therapy was only ∼30% in our current model, and subsequent stimulation with HBsAg/CpG-1826 vaccination did not further trigger anti-HBs transformation. We believe that the subtle differences in HBV tolerance extend among individual mice, and the insufficient immunomodulatory effects of existing IFN-α2, are the reasons for the low HBsAg seroconversion rate. Efforts to optimize the dosage and timing of both drugs may help improve the HBsAg seroconversion rate. Further development of other complementary immunomodulatory strategies is still necessary. Since our data indicates that there are immunosuppressive effects against B cells after long-term PEGIFNα treatment, for those recipients who did not achieve anti-HBs seroconversion, further PEGIFNα therapy may hardly improve the therapeutic outcomes but might instead add to immunosuppressive effects and lead to systemic toxicity. The issues of how to improve the efficacy and reduce the side effects of PEGIFNα have long been questioned. Our recent studies suggest that different subtypes of IFN-α have differential antiviral and immune-modulating activities. Compared with IFN-α2, subtypes of IFN-α with superior activity in activation of antiviral immune responses may improve the therapeutic efficacy.^46^ It is reasonable to anticipate that they will also be more effective when used in combination therapy, which is currently under study.

In conclusion, the present study evaluated the antiviral effects of combined RNAi and PEGIFNα treatment in chronic HBV-carrier IFNAR-hEC mice, showing synergistic antiviral efficacy, with a potent reduction of HBsAg of ∼4log10 IU/mL and a HBsAg seroconversion rate of ∼30%. The synergistic efficacy was mainly attributed to the ablation of high viral burden-induced immune tolerance by RNAi, the induction of global and virus-specific T cell and B cell responses by PEGIFNα, as well as the altered T cell and B cell communications by combined therapy.

## Contributions

Zhenghong Yuan, Jieliang Chen and Wenjing Zai conceived the project, designed the experiments and supervised the overall project. Wenjing Zai and Kongying Hu collected the samples and performed the virological experiments. Wenjing Zai and Mengying He analyzed the scRNA-seq data ad performed the flow cytometry experiments. Ziyang Song and Chen Luo helped performing the animal experiment. Minxiang Xie and Asha Ashuo provided the methods for T cell and B cell analysis. Wenjing Zai, Kongying Hu and Mengying He wrote the manuscript. All authors contributed to reading and editing the manuscript.

All authors have read and approved the article.

## Data sharing statement

Data are available on reasonable request. All data relevant to the study are included in the article or uploaded as supplementary information. Additional data are available upon reasonable request.

## Acknowledgements

The authors would like to thank the core facility of Shanghai Medical College, Fudan University for the technical platform and assistance. We would like to thank the Key Laboratory of Medical Molecular Virology (MOE/NHC/CAMS), Shanghai Frontiers Science Center of Pathogenic Microorganisms and Infection, School of Basic Medical Sciences, Shanghai Medical College, Fudan University and the faculty of the ABSL2 Laboratory Animal Facility for the technical platform and assistance, we sincerely thank Yuanfei Zhu, Yutang Li, and Qiliang Cai for their support in conducting animal experience.

